# Renal Cl-/H+ antiporter ClC-5 regulates collagen production and release in Dent Disease models

**DOI:** 10.1101/2023.04.21.537823

**Authors:** M. Duran, G. Ariceta, ME Semidey, C Castells-Esteve, B. Lu, A. Meseguer, G. Cantero-Recasens

## Abstract

Mutations in the Cl^-^/H^+^ antiporter ClC-5 cause Dent’s Disease 1 (DD1), a rare primary tubulopathy that eventually progresses to renal failure. In fact, even with normal kidney function, DD1 patients present renal tubulointerstitial fibrosis. However, the link between ClC-5 loss-of-function and renal fibrosis remains unclear. Here, we have shown that DD1 mice models lacking ClC-5 present higher renal collagen deposition and fibrosis. Accordingly, deletion of ClC-5 in human renal proximal tubule epithelial cells (*CLCN5* KD) recapitulates this effect. We have demonstrated that *CLCN5* KD causes an increase of collagen I (Col I) and IV (Col IV) intracellular levels by promoting their transcription through β-catenin pathway and impairing their lysosomal-mediated degradation. In addition, *CLCN5* KD cells release more Col I and IV at the extracellular space that form fibres with altered properties and resistance to removal compared to control cells. Altogether, we describe a new regulatory mechanism for collagens’ production and release by ClC-5, which is altered in DD1 and provides a better understanding of disease progression to renal fibrosis.

**SIGNIFICANCE STATEMENT:** Renal fibrosis is a common pathologic process occurring as consequence of chronic kidney injury and leading to renal dysfunction. Dent’s Disease is a rare renal pathology that progresses to chronic kidney disease and tubulointerstitial fibrosis. Interestingly, it is caused by mutations in a single gene called *CLCN5*, therefore it can help understanding the cellular mechanisms of renal fibrosis. Using cellular and mice models of the disease, we describe a mechanism linking *CLCN5* function, cell differentiation and regulation of collagen levels, major component of extracellular matrix and important player for renal fibrosis development. In conclusion, our results provide a link between *CLCN5* and altered collagen deposition, which could be relevant for other renal Fanconi syndrome related diseases also progressing to fibrosis.

## INTRODUCTION

Renal fibrosis is a common pathologic process occurring as consequence of chronic kidney injury, independently of the underlying disease aetiology (1). It is characterized by pathological deposition and accumulation of extracellular matrix (ECM) in the renal interstitium (2). Renal ECM is composed mainly of collagens, elastin and other glycoproteins to form the basal membrane. In normal conditions, ECM is in constant remodelling state, with an equilibrium between degradation of old or damaged molecules and replacement by new ones. In fibrosis, however, this is altered with increased production and reduced degradation, leading to excessive ECM deposition (3). Indeed, collagen turnover is a key event for ECM remodelling and impairment of this process leads to fibrosis initiation and progression (4). Interestingly, up to 30% of newly synthesized collagen is sent for intracellular degradation (5,6) as a mechanism to prevent defective collagen’s secretion and therefore preserving the quality of ECM. In addition, collagens type I, II, III, IV, V, VI, VII, and XV levels are increased in renal fibrosis (1,3).

ClC-5 (Cl^-^/H^+^ exchange transporter 5), encoded by *CLCN5* gene, is mainly expressed in renal epithelial cells of proximal tubule (PTC) (7). It controls endosomal acidification by cooperating with the V-ATPase, which affects protein recycling to the plasma membrane (e.g. endocytic receptors) (8). Accordingly, ClC-5 loss-of-function leads to alterations in endocytic activity (9) and delays protein progression to lysosomes thereby affecting the functioning of PTCs (8). In fact, mutations on *CLCN5* cause Dent’s disease type 1 (DD1), a rare X-linked renal proximal tubulopathy characterized by hypercalciuria and low molecular weight proteinuria (10,11). In addition, some patients may show an incomplete or complete renal Fanconi Syndrome (12). DD1 progresses to chronic kidney disease and finally to renal failure requiring renal replacement therapy between the thirties-fifties in up to 80% of patients (13). Importantly, several DD1 patients present tubulointerstitial fibrosis even with normal renal function (14). Besides, kidney biopsies from DD1 patients demonstrated progressive interstitial fibrosis associated with glomerular sclerosis overtime, in parallel to glomerular filtration decline (15). Interestingly, lack of ClC-5 has been shown to lead to epithelial cell dedifferentiation and decrease in endocytic receptors in both mice and cell models (16–19). However, although ClC-5 loss-of-function is the cause of DD1 and its role as Cl^-^/H^+^ exchanger is well studied, the link between ClC-5 and renal fibrosis remains unknown.

Here, we show in cellular and mice DD1 models that ClC-5 modulates intracellular and extracellular collagen levels by controlling epithelial cell differentiation and lysosomal collagen degradation, thus contributing to renal fibrosis.

## RESULTS

### 1. Mice lacking *Clcn5* present a renal fibrotic phenotype with increased Col IV deposition

*Clcn5* knockout mice (*Clcn5*^*-/-*^ mice) reproduce human DD1 pathophysiology, presenting low molecular weight proteinuria and reduced uptake of filtered proteins at proximal tubule (16,17). However, study of fibrosis in these mice has not been addressed, thus, we proceeded to evaluate it in kidneys from *Clcn5*^*+/+*^, *Clcn5*^*+/-*^, *Clcn5*^*-/-*^ mice. To facilitate the analysis and taking into account that heterozygous mice also present DD1 phenotype (20), *Clcn5*^*+/-*^ *and Clcn5*^*-/-*^ mice were grouped together. Haematoxylin and eosin (H/E) staining showed no obvious histological differences in glomeruli structure, tubule formation or inflammatory infiltrate between conditions (**Figure 1A, Supplementary Table 1**). We then stained the kidney slices with Sirius Red/Fast Green to determine total levels of collagens. This staining revealed that *Clcn5*^*+/-*^ *and Clcn5*^*-/-*^ mice present a thicker basal membrane than *Clcn5*^*+/+*^ mice (1.39 μm *vs*. 0.76 μm, p<0.05) (**Figure 1A-B**). Specific staining of Collagen type IV (Col IV), which is increased in renal fibrosis, revealed higher extracellular deposition of Col IV in mice lacking *Clcn5* (*Clcn5*^*+/-*^ *and Clcn5*^*-/-*^) compared to *Clcn5*^*+/+*^ mice (1.14 μm *vs*. 0.81 μm, p<0.01) (**Figure 1A and 1C**). Accordingly, *Col4a1* (Col IV) mRNA levels were also increased in kidneys from *Clcn5*^*+/-*^ *and Clcn5*^*-/-*^ mice (1.6-fold increase, p<0.05) (**Figure 1D**). Finally, global assessment confirmed that *Clcn5*^*+/-*^ and *Clcn5*^*-/-*^ mice present higher level of renal fibrosis than *Clcn5*^*+/+*^ mice (Fibrosis: 66.6% *vs*. 0%, p<0.01) (**Figure 1E**).

**Figure 1.**
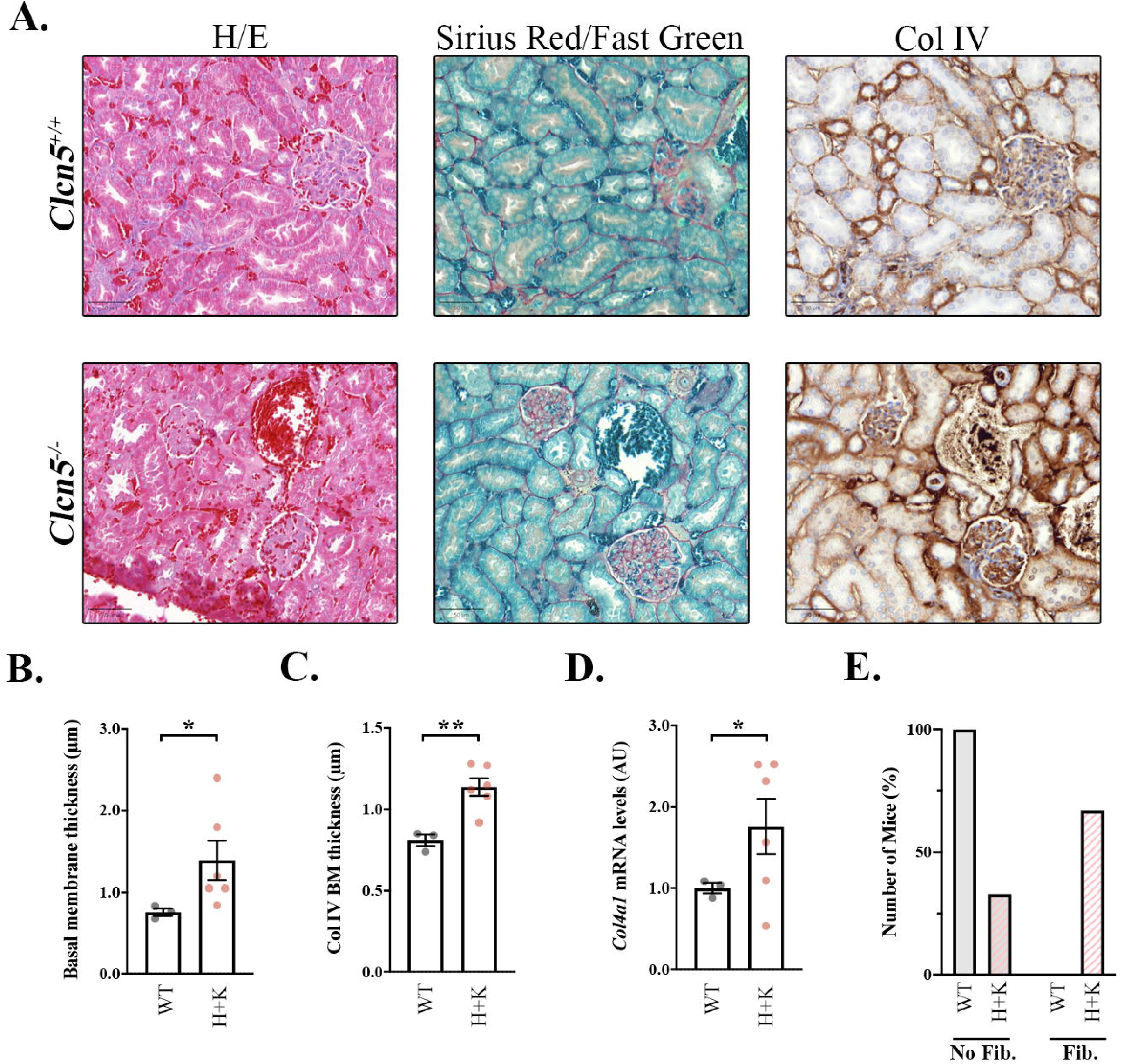
*Clcn5*^*+/-*^ and *Clcn5*^*-/-*^ mice show increased basal membrane thickness. **(A)** Representative kidney slices of *Clcn5*^*+/+*^ (upper panel) or *Clcn5*^*-/-*^ (lower panel) mice stained with Haematoxylin/Eosin (H/E), Sirius Red/Fast Green (Red: all collagens, green: non-collagenous proteins) or against Collagen type IV (brown, Col IV). Scale bar = 50 μm. **(B**,**C)** Quantification of the basal membrane thickness in the kidneys stained with Sirius Red/Fast Green **(B)** or against Col IV **(C)** of *Clcn5*^*+/+*^ (WT, grey dots) or *Clcn5*^*+/-*^ and *Clcn5*^*-/-*^ mice (H+K, pink dots). Average values ± SEM are plotted as scatter plot with bar graph. The y-axis represents the thickness of the basal membrane in μm. Statistical significance was determined using two-tailed unpaired t-test with Welch’s correction. **(D)** *Col4a1* (Col IV) mRNA levels from kidneys of *Clcn5*^*+/+*^ (WT, grey dots) or *Clcn5*^*+/-*^ and *Clcn5*^*-/-*^ mice (H+K, pink dots) normalised by *Hprt1* levels. Average values ± SEM are plotted as scatter plot with bar graph. Statistical significance was determined using one-tailed unpaired t-test with Welch’s correction. **(E)** Percentage of *Clcn5*^*+/+*^ (WT) or *Clcn5*^*+/-*^ and *Clcn5*^*-/-*^ mice (H+K) not presenting any evidence of fibrosis (No Fib.) or showing some level of fibrosis (Fib.). Abbreviations: H/E: Haematoxylin/Eosin staining, Col IV: Collagen type IV, WT: *Clcn5*^*+/+*^ mice, H+K: *Clcn5*^*+/-*^ and *Clcn5*^*-/-*^ mice, BM: Basal Membrane. * p<0.05, ** p<0.01

### 2. Depletion of ClC-5 alters MUC1 levels

To further study ClC-5 role on collagen production and renal fibrosis, we used previously generated renal proximal tubule epithelial cell lines (RPTEC/TERT1) depleted of endogenous *CLCN5 (CLCN5* KD) or overexpressing ClC-5 WT rescue form (rWT) (18). These cells lacking a functional ClC-5 present several characteristics of epithelial dedifferentiation (18), which we postulate that promotes an increase in collagens’ production that could explain renal fibrosis progression in DD1.

First, analysis of *CLCN5*/ClC-5 levels confirmed >90% reduction in *CLCN5* KD cells and complete expression recovery in rWT cells compared to control cells (**Supplementary Figure 1A-B**). Confirming previous results (18), *CLCN5* depleted cells showed strong reduction in E-cadherin protein levels compared to control and rWT cells (>90% decrease, p<0.05) (**Figure 2A**). We also analysed the levels of mucin-1 (MUC1), a relevant cell polarization and differentiation marker (25), and one of the most affected genes by ClC-5 loss-of-function as shown by microarray data (18). Accordingly, *CLCN5* deletion caused a decrease in MUC1 protein levels compared to control cells, which was partially recovered by expression of ClC-5 rWT (**Figure 2B**). Confocal immunofluorescence analyses showed that *CLCN5* deletion not only caused a reduction of MUC1 levels, but also affected its localization at the plasma membrane (PM). This defect in MUC1 trafficking was almost completely restored by expression of ClC-5 rWT (**Figure 2C**).

**Figure 2.**
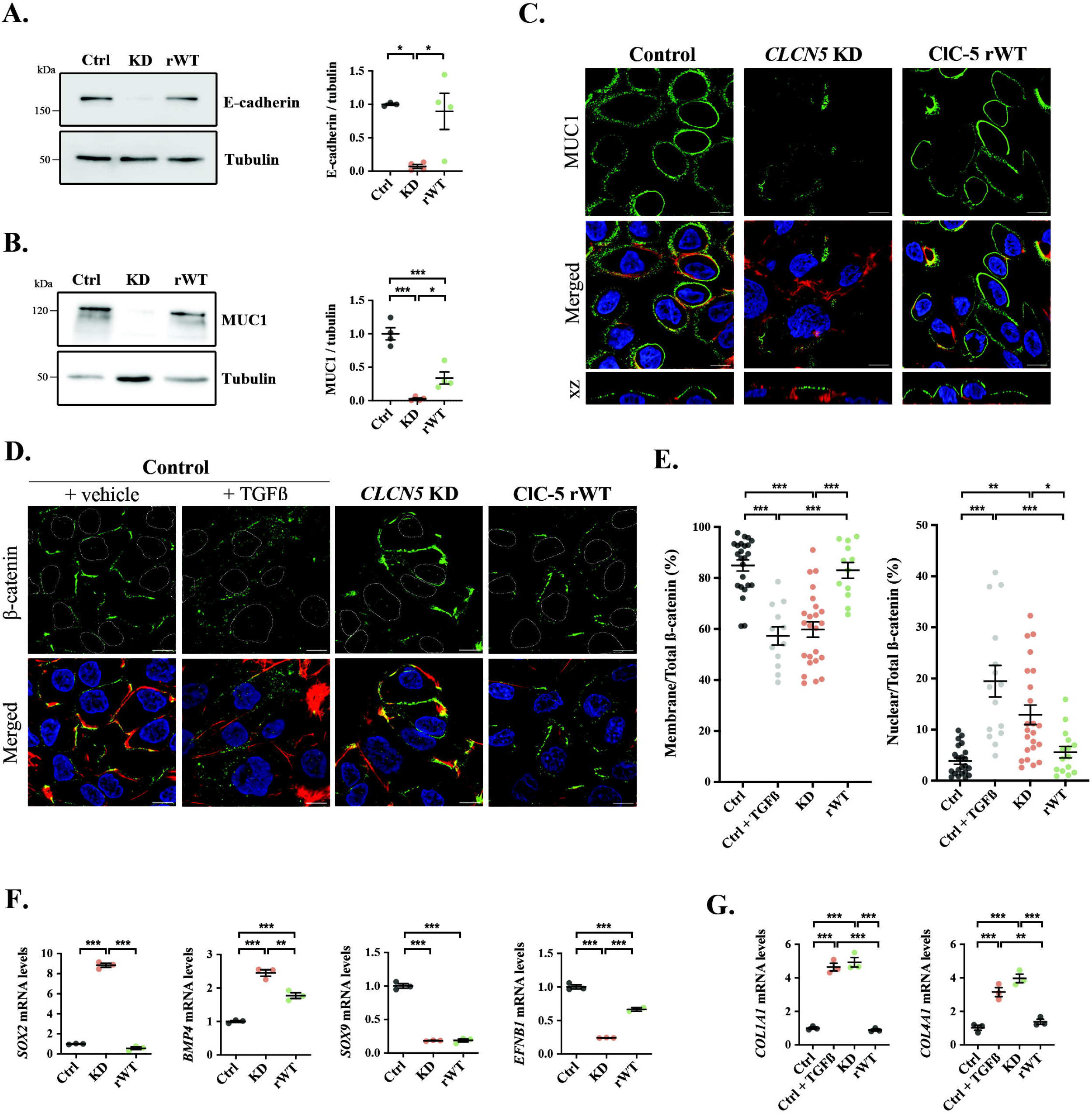
*CLCN5* KD cells present decreased epithelial marker levels and higher β-catenin nuclear translocation. **(A)** E-cadherin protein levels were analysed by western blot in cell lysates from control, *CLCN5* KD and ClC-5 rWT cells. Tubulin was used as a loading control. Right graph shows the quantification of E-cadherin levels (N>3). **(B)** MUC1 protein levels were analysed by western blot in cell lysates from control, *CLCN5* KD and ClC-5 rWT cells. Tubulin was used as a loading control. Right graph shows the quantification of MUC1 levels (N>3). **(C)** Immunofluorescence z-stack single planes of control, *CLCN5* KD, and ClC-5 rWT cells stained with anti-MUC1 antibody (green), phalloidin (red), and Hoechst 33342 (blue). Scale bars: 5 μm. **(D)** Immunofluorescence single planes of control (treated with vehicle or TGFβ as positive control), *CLCN5* KD, and ClC-5 rWT cells stained with anti-β-catenin (Green), phalloidin (red), and Hoechst 33342 (blue). Nuclei perimeter is demarcated with dotted white lines to visualise β-catenin nuclear translocation. Scale bars: 5 μm. **(E)** β-catenin levels at the membrane (left graph) or at the nucleus (right graph) normalised to total levels were quantified from immunofluorescence images of control + vehicle (black dots), control + TGFβ (grey dots), *CLCN5* KD (pink dots) and ClC-5 rWT (green dots) cells. **(F)** *SOX2, BMP4, SOX9* and *EFNB1* mRNA levels (normalised to *TBP* levels) from control (black dots), *CLCN5* KD (pink dots) and ClC-5 rWT (green dots) cells. **(G)** *COL1A1* (Col I) and *COL4A1* (Col IV) mRNA levels (normalised to *HPRT1* levels) from control (black dots), *CLCN5* KD (pink dots) and ClC-5 rWT (green dots) cells. Abbreviations: Ctrl: Control cells, KD: *CLCN5* KD cells, rWT: ClC-5 rWT cells. Statistical significance was determined using one-way ANOVA followed by Tukey’s post hoc test. * p<0.05, ** p<0.01, *** p<0.001.

### 3. ClC-5 deletion activates β-catenin pathway

E-cadherin can form a complex with β-catenin, and its loss can promote β-catenin release from PM and translocation to the nucleus, facilitating epithelial-mesenchymal transition (EMT), promoting collagen transcription and leading to renal fibrosis (26–28). Furthermore, β-catenin can also bind to MUC1 (29). Thus, since ClC-5 deletion reduces both E-cadherin/MUC1 levels, we tested β-catenin localization. Our results revealed that β-catenin is mostly localized at PM in control cells and is translocated to the nucleus after stimulation by the pathway activator TGF-β (30) (5-fold increase, p<0.01). Importantly, β-catenin was also found at the nucleus in *CLCN5* KD cells, which was prevented by ClC-5 rWT expression (KD: 3.3-fold increase, p<0.05; rWT: 1.4-fold increase, n.s., compared to control cells) (**Figure 2D-E**). Next, we studied whether this translocation promoted the expression of known β-catenin target genes (31). Analysis of the public data from DD1 cellular models microarrays (18) showed that several of β-catenin target genes are also modulated by *CLCN5* deletion (**Supplementary Table 2**). Four representative genes *(BMP4, SOX2, SOX9* and *EFNB1*) were selected to validate these data. Accordingly, *SOX2* and *BMP4* were upregulated (8- and 2.5-fold increase, respectively), while *SOX9* and *EFNB1* were downregulated (80% and 75% reduction, respectively) in *CLCN5* KD cells. ClC-5 rWT expression partially recovered *BMP4* and *EFNB1* levels, suggesting that other pathways rather than β-catenin may be regulating the expression of the other genes (**Figure 2F**).

### 4. Lack of ClC-5 enhances Col I and IV production and release

β-catenin nuclear translocation promotes collagen transcription (32,33), which would lead to EMT and renal fibrosis. In accordance, our data revealed that Collagen type I (Col I) and Collagen type IV (Col IV) mRNA levels, both increased in renal fibrosis, were higher in *CLCN5* KD cells than in control or rWT cells (4.9 and 4.0-fold increase, respectively, p<0.01). A similar increment was detected after TGF-β stimulation, suggesting β-catenin involvement in the control of collagens expression by ClC-5 (**Figure 2G**). To sum up, our data reveal that ClC-5 depletion promotes Col I/IV transcription most likely through β-catenin translocation.

Uncontrolled collagens’ release and accumulation at tubulointerstitial space is characteristic of renal fibrosis, a process occurring in most DD1 patients (14). Our data have confirmed that Col I/IV mRNA levels are increased in cells depleted of *CLCN5*. However, does it translate to augmented Col I/IV protein levels, which could lead to renal fibrosis? Our results revealed that ClC-5 depletion caused a strong increase in intracellular Col IV and Col I protein levels compared to control or rWT cells (10-fold and 4-fold increase, respectively) (**Figure 3A-E**). Extracellular levels of Col IV and Col I were also elevated in *CLCN5* KD cells compared to control or rWT cells (4 and 6-fold increase, respectively) (**Figure 3A-E**). Immunofluorescence images confirmed a massive increase of intracellular Col IV and Col I levels in *CLCN5* KD cells compared to control and rWT cells (**Figure 3C and 3F**, respectively).

**Figure 3.**
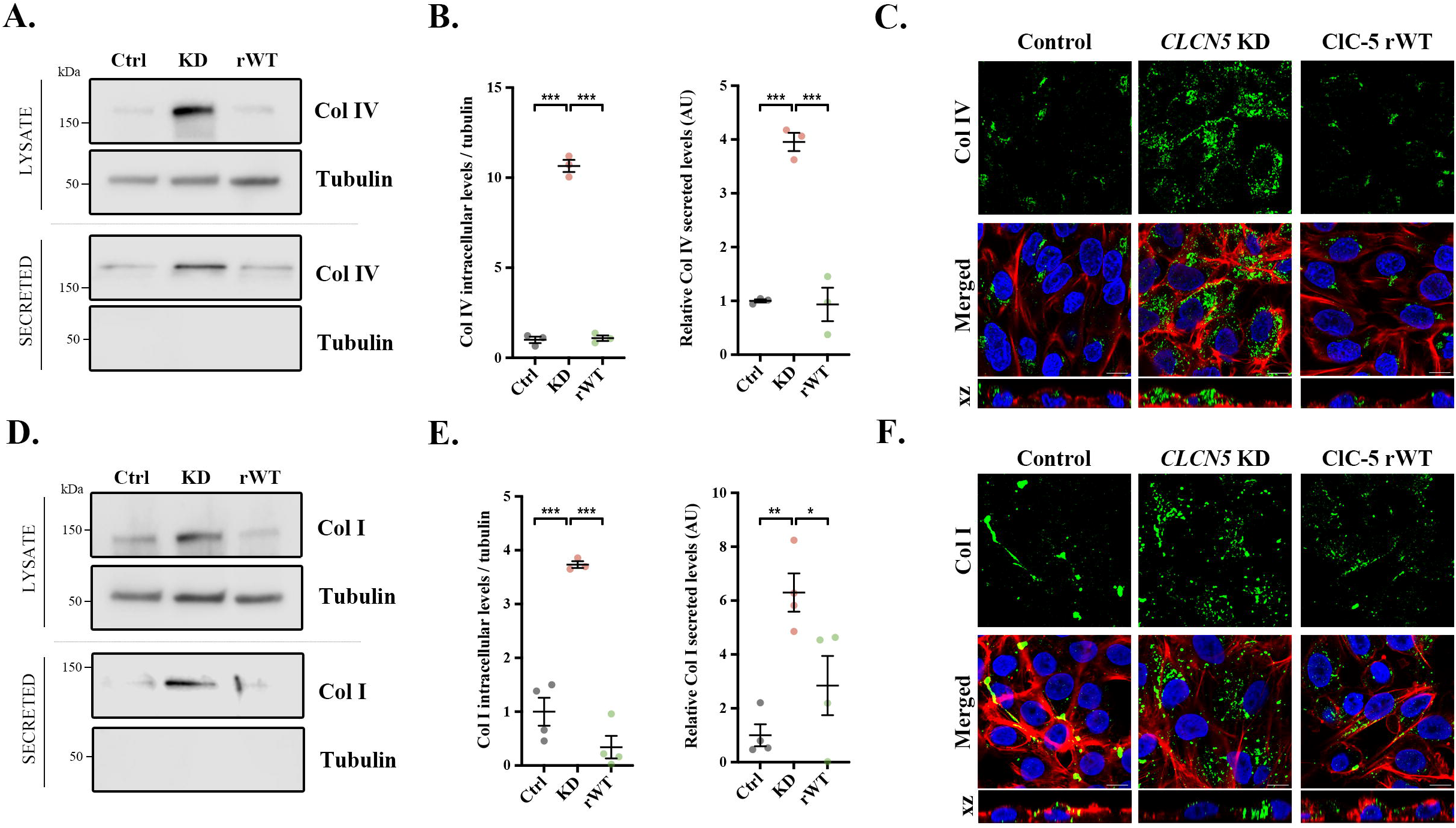
Collagen type I and type IV intracellular and extracellular levels are increased in *CLCN5* KD cells. **(A, D)** Collagen type IV (Col IV) **(A)** or Collagen type I (Col I) **(D)** protein levels in the lysates or secreted media of control (Ctrl), *CLCN5* KD (KD) and ClC-5 rWT (rWT) cells. Tubulin was used as a loading control in the lysates, and as a control of lysate contamination in secreted media. **(B, E)** Quantification of Col IV **(B)** or Col I **(E)** protein intracellular levels (left graph) or secreted levels (right graph) of control (Ctrl, black dots), *CLCN5* KD (KD, pink dots) and ClC-5 rWT (rWT, green dots) cells (N = 3). **(C, F)** Representative immunofluorescence z-stack single planes of control, *CLCN5* KD, and ClC-5 rWT cells stained with anti-Col IV **(C)** or anti-Col I **(F)** antibody (green), phalloidin (red), and Hoechst 33342 (blue). Scale bars: 5 μm. Abbreviations: Col I: Collagen type I, Col IV: Collagen type IV, Ctrl: Control cells, KD: *CLCN5* KD cells, rWT: ClC-5 rWT cells. Statistical significance was determined using one-way ANOVA followed by Tukey’s post hoc test. * p<0.05, ** p<0.01, *** p<0.001.

### 5. ClC-5 deletion impairs Col I/IV degradation

Enhanced transcription can partially explain the increment of Col I/IV intracellular levels by ClC-5 depletion. Nonetheless, considering that 30% of newly synthesised collagen is normally degraded (34), and since ClC-5 is a key component for endolysosomal system, we postulate that the increase in collagens’ intracellular levels could also be caused by impairment of their degradation. Study of Col I and IV half-life revealed that it is extremely delayed in *CLCN5* KD cells compared to control cells (11.2 vs. 2.2 hours and 13.1 vs. 1.75 hours, respectively). Overexpression of ClC-5 rWT completely rescues this phenotype (Col I: 3.3 h and Col IV: 2.5 h) (**Figure 4A-B**). Altogether, our data demonstrate that collagens’ degradation is affected by ClC-5 deletion.

**Figure 4.**
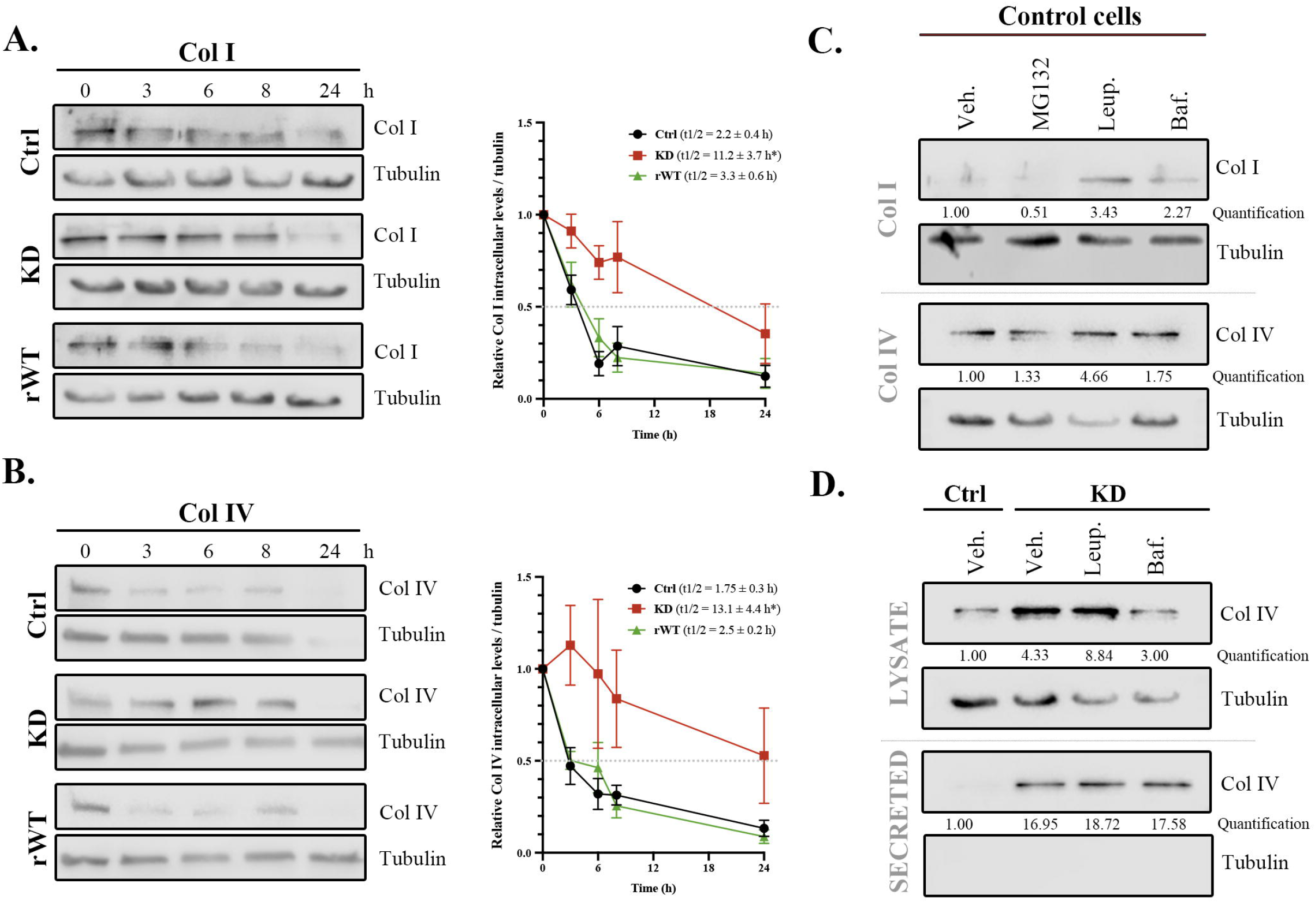
Collagen lysosomal degradation is impaired in cells lacking *CLCN5*. **(A, B)** Representative blots of cycloheximide chase assay for Col I **(A)** and Col IV **(B)** protein levels over time (0, 3, 6, 8 and 24 h) of control (Ctrl), *CLCN5* KD (KD) and ClC-5 rWT (rWT) cells. Quantification of respective protein levels for each time point normalised to tubulin and relative to time = 0 h is shown in the graph. Half-life of Col I or Col IV is shown for each cell line (N > 3). Statistical significance was determined using one-way ANOVA followed by Tukey’s post hoc test. * p<0.05 (*CLCN5* KD compared to Control and ClC-5 rWT cells). **(C)** Western blots of Col I (Upper lanes) or Col IV (lower lanes) from lysates of control cells treated with vehicle (Veh.), MG132, Leupeptin (Leup.) or bafilomycin (Baf.). Tubulin was used as a loading control. Quantification of Col I and Col IV are included. **(D)** Col IV intracellular (lysate) or extracellular (secreted) levels from control cells (Ctrl) or *CLCN5* KD cells (KD) treated with vehicle (Veh.), Leupeptin (Leup.) or Bafilomycin (Baf.). Tubulin was used as a loading control. Quantification of Col I and Col IV are included. Abbreviations: Col I: Collagen type I, Col IV: Collagen type IV, Ctrl: Control cells, KD: *CLCN5* KD cells, rWT: ClC-5 rWT cells, Veh: Vehicle, Leup: Leupeptin, Baf: Bafilomycin.

To understand whether ClC-5 depletion affects lysosomal or proteasomal degradation, we studied how collagens are degraded in renal proximal tubule epithelial cells (RPTEC) using different inhibitors (MG132, Leupeptin and Bafilomycin-A1). Our results showed that only inhibitors of lysosomes caused Col I and IV intracellular accumulation compared to vehicle (Leupeptin or Bafilomycin-A1: Col I, 3.4- and 2.2-fold increase; Col IV: 4.7- and 1.8-fold increase, respectively) (**Figure 4C**). Next, to confirm that lysosomal degradation of collagens is impaired due to ClC-5 deletion, we treated *CLCN5* KD cells with lysosomal inhibitors and analysed Col IV levels. We found that only Leupeptin (inhibitor of lysosomal enzymes), but not Bafilomycin-A1 (inhibitor of v-ATPase), further increased Col IV intracellular levels in *CLCN5* KD cells (**Figure 4D**). These data strongly suggest that ClC-5 impairs collagens degradation by altering lysosomal acidification. Importantly, none of the treatments affected the increment in extracellular collagen levels produced by ClC-5 deletion (**Figure 4D**). Next, we analysed whether Col I/IV were accumulated in lysosomes of *CLCN5* KD cells. Our results showed major colocalization of Col IV with LAMP1 (marker of lysosomes) in *CLCN5* KD cells compared to control or rWT cells (12 and 15-fold increase, respectively; p<0.05) (**Figure 5A, 5C**), consistent with impaired lysosomal degradation. Similarly, Col I was also more frequently found in the lysosomes of *CLCN5* KD cells than control cells (30-fold increase, p<0.05), although in this case expression of ClC-5 rWT was not sufficient to rescue the phenotype (**Figure 5B, 5C**). Altogether, our data revealed that Col I/IV are mainly degraded through lysosomes in RPTEC cells, a process impaired in cells lacking ClC-5, which also causes collagens to accumulate in the lysosomes.

**Figure 5.**
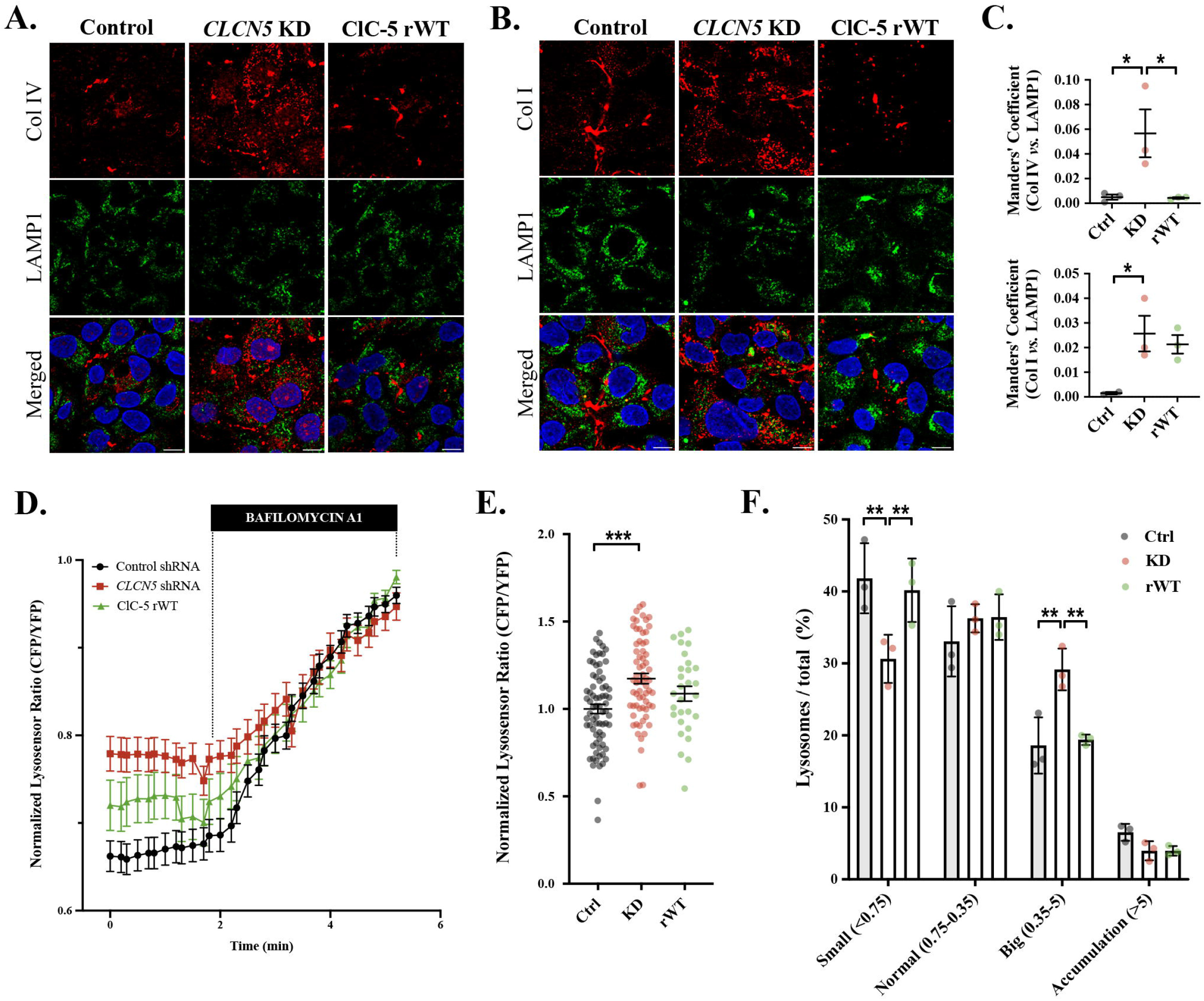
Lysosomes from *CLCN5* KD cells are less acidic and larger. **(A, B)** Representative immunofluorescence z-stack single-plane images of control, *CLCN5* KD, and ClC-5 rWT cells stained with anti-Col IV **(A)** or anti-Col I **(B)** antibody (red), LAMP1 (green), and Hoechst 33342 (blue). Scale bars: 5 μm. **(C)** Colocalization between Col IV (upper graph) or Col I (lower graph) and LAMP1 for different cell lines (Control, *CLCN5* KD and ClC-5 rWT) were calculated from immunofluorescence images by Manders’ coefficient using FIJI. Average values ± SEM are plotted as scatter plot with bar graph. The y-axis represents Manders’ coefficient of the fraction of Col I or IV overlapping with LAMP1. **(D)** Time course of CFP/YPF LysoSensor™ Yellow/blue DND-160 dye ratio in control (Control shRNA, black), *CLCN5* KD (*CLCN5* shRNA, red) and ClC-5 rWT cells. After 2 minutes of basal, bafilomycin was added as a normalization control. **(E)** Quantification of the normalised basal CFP/YFP LysoSensor™ Yellow/blue DND-160 dye ratio. Each dot represents one cell. **(F)** Percentage of the populations of lysosomes of control (Ctrl), *CLCN5* KD (KD) and ClC-5 rWT (rWT) cells classified by volume: Small (<0.75 μm^3^), Normal (between 0.75-0.35 μm^3^), Big (between 0.35-5 μm^3^) and accumulations (>5 μm^3^). Abbreviations: Col I: Collagen type I, Col IV: Collagen type IV, Ctrl: Control cells, KD: *CLCN5* KD cells, rWT: ClC-5 rWT cells. Statistical significance was determined using one-way ANOVA followed by Tukey’s post hoc test. * p<0.05, ** p<0.01, *** p<0.001.

### 6. CLCN5 depletion affects lysosomal acidification

Other authors have shown that ClC-5 has a role in endosomal acidification (8). However, is ClC-5 also affecting lysosomal pH in RPTEC cells? To test this possibility, we measured the luminal lysosomal pH using the LysoSensor™ Yellow/blue DND-160 dye. Colocalization of LysoSensor™ with LAMP1 showed that after 5 minutes, this dye is mostly found in the lysosomes (**(Supplementary figure 2A**). Thus, all subsequent experiments were performed after 5 minutes incubation to ensure LysoSensor™ is at lysosomes. Our data showed that in basal conditions lysosomes’ luminal pH of *CLCN5* KD cells is more basic than those of control cells (20% increase, p<0.01), and is partially recovered by expression of ClC-5 rWT (**Figure 5D-E**). Next, we studied whether localization, size or number of lysosomes were altered in cells lacking ClC-5. Immunofluorescence studies using anti-LAMP1 antibody and anti-KDEL antibody revealed a higher colocalization between both markers in *CLCN5* KD cells than control or rWT cells (2-fold increase, p<0.05) (**Supplementary figure 2B-C**), supporting the defect on lysosomal collagens’ degradation in cells depleted of ClC-5. In addition, our results showed increased percentage of big lysosomes in *CLCN5* KD cells compared to control or rWT cells (29.2±1.7%, 18.6±2.3% and 19.4±0.4%, respectively) (**Figure 5F**). Volume measurement also showed that lysosomes from *CLCN5* KD cells were bigger than control or rWT cells (Median volume of KD: 0.18 μm^3^, control: 0.11 μm^3^, rWT: 0.10 μm^3^, p<0.01) (**Supplementary Figure 2D**). These data confirm that ClC-5 depletion impairs lysosomal acidification, leading to defects on collagen degradation and causing an increase in Col I/IV intracellular levels.

### 7. Col I/IV extracellular levels are increased in *CLCN5* KD cells

Our data have provided a possible mechanism explaining how ClC-5 loss-of-function can enhance synthesis and impair degradation causing intracellular collagens’ accumulation. However, what is the connection between ClC-5 and the increase in collagens’ extracellular levels observed in *CLCN5* KD cells? Chemical inhibition of lysosomes, which causes intracellular accumulation of Col I/IV, also produces an increase of extracellular collagen levels in control cells, but not in *CLCN5* KD cells (**Supplementary Figure 3A** and **Supplementary Figure 4D**). TGF-β treatment that promotes Col I/IV transcription, also causes an increase of their levels at external medium (**Figure 3B**). Therefore, the increment in extracellular Col I/IV levels is possibly due to increased intracellular levels. But, considering ClC-5 deletion induces a loss of cell polarization, where are collagens localized extracellularly and how are they arranged in these cells?

To answer these questions, Col I/IV fibres were analysed before or after extensive washing. Surprisingly, the phenotype observed in *CLCN5* KD cells was different between Col I and Col IV. On one hand, before washing *CLCN5* KD cells presented higher amounts of Col I large fibres than control and rWT cells (11.7, 2.7 and 3.3 fibres/field, respectively; p<0.05). After extensive washing, many Col I fibres were still detected in control and rWT cells (7.3 and 5.0 fibres/field, respectively; n.s. compared to non-washed cells), while they were virtually absent in *CLCN5* KD cells (1.3 fibres/field, 90% reduction compared to non-washed cells; p<0.05) (**Figure 6A-B**). Measurement of Col I fibres’ area also confirmed large fibres were removed after washing (**Supplementary Figure 3C**). On the other hand, although there was a tendency of more Col IV large fibres in non-washed *CLCN5* KD cells compared to control or rWT cells (14.3, 7.0, 4.0 fibres/field, respectively; n.s.), most of these fibres remained after washing only *CLCN5* KD, but not in control or rWT cells (12.7, 3.3, 0.0 fibres/field, respectively; p<0.05) (**Figure 6C-D**). As showed in Figure 6C, Col IV fibres were located in the intercellular space when ClC-5 was deleted, which could explain their resistance to washing and be related to renal fibrosis phenotype (**Figure 6C**). Quantification of Col IV fibres’ area also showed that large fibres are not removed after washing (**Supplementary Figure 3D**).

**Figure 6.**
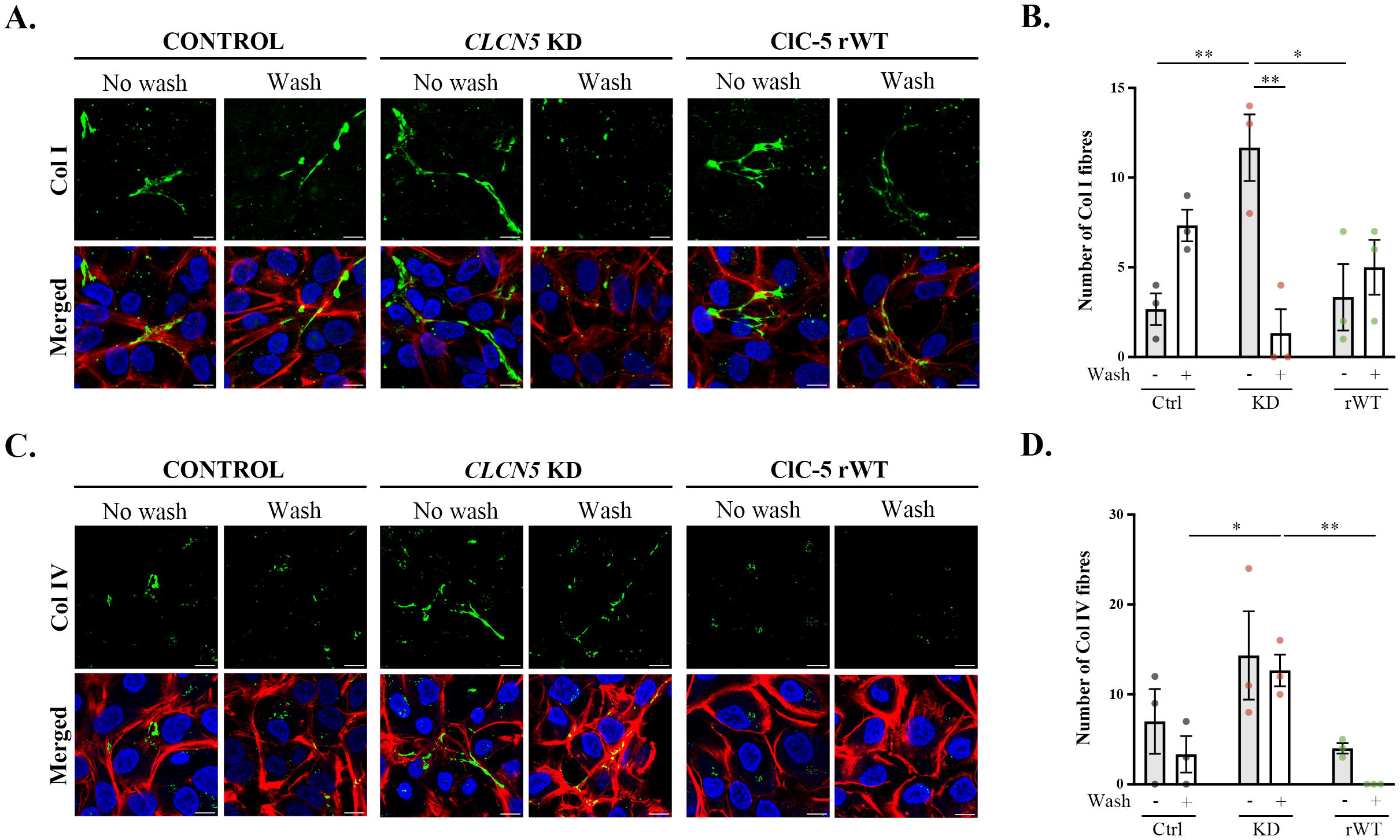
*CLCN5* KD cells accumulate more collagens extracellularly. **(A, C)** Control, *CLCN5* KD and ClC-5 rWT cells were processed for immunofluorescence before (No wash) or after extensive washing (Wash). Cells were stained with anti-Col I **(A)** or anti-Col IV **(C)** antibodies (Green), phalloidin (Red) and Hoechst 33342 (Blue). Scale bars = 5 μm.. **(B)** Quantification of the number of Col I fibres (defined as those Col I particles with an area > 2 μm^2^, perimeter > 10μm) in control (Ctrl), *CLCN5* KD (KD) and ClC-5 rWT (rWT) cells before and after washing. **(C)** Quantification of the number of Col IV fibres (defined as those Col I particles with an area > 1.5 μm^2^, perimeter > 10μm) in control (Ctrl), *CLCN5* KD (KD) and ClC-5 rWT (rWT) cells before and after washing. Abbreviations: Col I: Collagen type I, Col IV: Collagen type IV, Ctrl: Control cells, KD: *CLCN5* KD cells, rWT: ClC-5 rWT cells. Statistical significance was determined using one-way ANOVA followed by Tukey’s post hoc test. * p<0.05.

## DISCUSSION

Mutations on *CLCN5*, the genetic cause of Dent’s Disease type 1 (DD1), lead to proximal tubule epithelial cells dedifferentiation and dysfunction (8,18). Importantly, most DD1 patients will develop renal fibrosis with age and progress to renal failure (35). Renal fibrosis is characterized by pathological accumulation of extracellular matrix (ECM) in the renal interstitium, which is mainly composed of collagens, being Col I and Col IV specifically increased in renal fibrosis (1–3). However, how can ClC-5, a Cl^-^/H^+^ antiporter located at endosomes and PM, have a role on the production and release of collagens?

Here, we provide a plausible novel mechanism linking ClC-5 with collagens and renal fibrosis. We propose that ClC-5 modulates Col I/IV intracellular and extracellular levels by controlling their synthesis via β-catenin pathway and degradation by lysosomes. In the absence of ClC-5, there is a dysregulation of this system that ends in collagens’ overproduction and release, which will lead to renal fibrosis, mimicking what is observed in up to 60% of DD1 patients’ kidney biopsies (15).

Our data show that *Clcn5* KO (*Clcn5*^*+/-*^) and heterozygous (*Clcn5*^*-/-*^) mice already present evidences of renal fibrosis at 3-months of age, reproducing this DD1 feature. One of the main findings is the thicknessing of the basal membranes in *Clcn5*^*+/*^ and *Clcn5*^*-/-*^ mice, which is also clearly observed by specific Col IV staining, a major element of renal ECM. Accordingly, other authors have shown that ClC-5 overexpression in unilateral ureteral occlusion (UUO) mice cause a decrease of Col IV levels and ameliorates UOO-induced renal injury (36). This increase in collagen levels was also reproduced in RPTEC cells depleted of *CLCN5*, which showed a dramatic intracellular Col I/IV accumulation.

Using DD1 cellular models, we have demonstrated that this increase was due to enhanced synthesis and impaired degradation. First, we have found that ClC-5 deletion promotes downregulation of MUC1/E-cadherin and β-catenin nuclear translocation, which leads to transcription of several EMT-related genes. Interestingly, both MUC1/E-cadherin can anchor β-catenin at PM, preventing its function (29,37). Therefore, a reduction in their levels would permit β-catenin nuclear translocation. Besides, MUC1/E-cadherin levels are reduced after β-catenin nuclear translocation (28,29), indicating a feedback loop that further decreases epithelial markers’ expression and enhances β-catenin translocation. Surprisingly, studies on cancer cells have shown the opposite effect of ClC-5 on β-catenin (38). In cancer cells, ClC-5 inhibition reduces β-catenin levels and nuclear translocation, leading to apoptosis and having anti-tumoral effect. We speculate these differences could be because cancer cells start overexpressing ClC-5, which may enhance acidification of endolysosomal pathway and promote tumour resistance by degrading potential markers or receptors. Second, we have found that Col I/IV are mostly degraded lysosomally in RPTEC cells, a process that is altered in *CLCN5* KD cells. In fact, and contrarily to mainstream dogma (8), our results show that *CLCN5* depletion does affect the lysosomal pH. Another possibility we cannot fully discard is that by affecting the endolysosomal system pH, deletion of *CLCN5* is not directly affecting collagens’ degradation, but their targeting to lysosomes. Our results, indeed, also support this hypothesis as little percentage of total collagens (although higher than in control conditions) is located in lysosomes compared to the amount found in endoplasmic reticulum of *CLCN5* KD cells. To sum up, we have demonstrated that 1) collagens are lysosomally degraded in RPTE cells and 2) ClC-5 deletion increases intracellular collagen levels by upregulating their expression and impairing their degradation.

This upregulation of Col I/IV intracellular levels caused an increase of extracellular levels, although the secretion rate remained unaffected by *CLCN5* KD. Interestingly, secreted Col I/IV by *CLCN5* KD or control cells form fibres with different resistance to washing. While Col IV is stacked between cells in absence of ClC-5; Col I forms big fibres that are easily removed from KD cells after extensive washing. This suggests that on one hand, lack of epithelial markers causes a loss of cell polarity, allowing collagens to be secreted not only basolaterally. This could also partially explain the increase in collagens’ extracellular levels, and why Col IV is detected in the intercellular space instead of being at the basal membrane. In the case of Col I, which is not usually directly attached to PM, we postulate that anchoring proteins that make Col I fibres resistant to washing are specific of the basolateral site. Thus, Col I secreted apically cannot bind any anchoring protein and is more sensitive to removal by washing.

## CONCLUSIONS

Our data indicates that normal RPTEC cells tightly control collagens’ production and release, to prevent dysregulated accumulation that could be detrimental for renal function and lead to renal fibrosis. Based on our results, we propose a model where ClC-5 is a key regulator of collagens synthesis and secretion by modulating the acidification of the endolysosomal system, which impacts protein intracellular trafficking and lysosomal degradation (**Figure 7**). Altogether, our data provides a new mechanism for ClC-5 role on renal fibrosis by regulating Col I/IV levels. In addition, our results may be also relevant for other renal Fanconi Syndrome related diseases also progressing to renal fibrosis.

**Figure 7.**
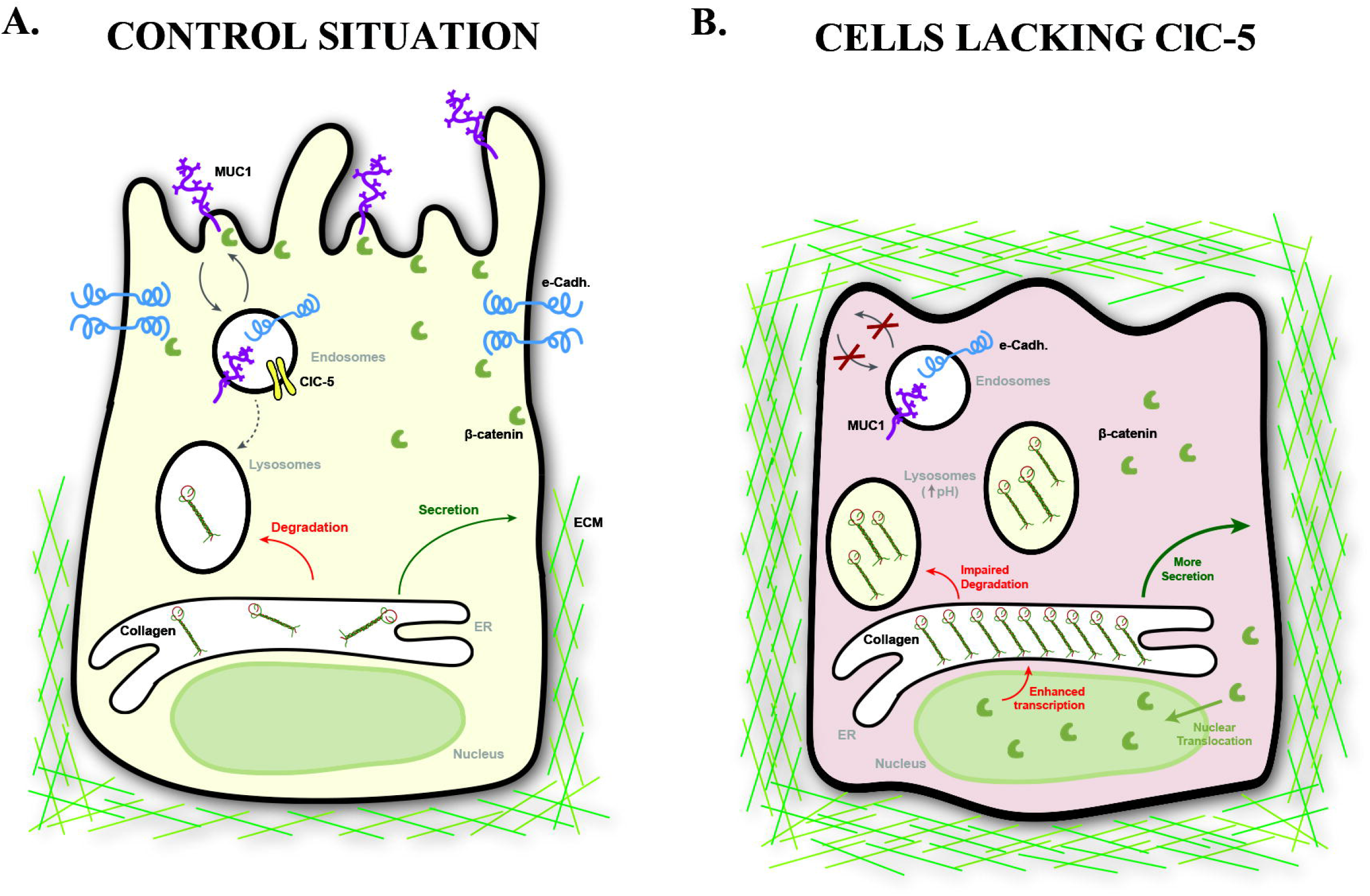
Model for ClC-5 role on renal fibrosis in DD1. **(A)** In normal situation, ClC-5 maintains the trafficking of epithelial markers to PM, which sequesters β-catenin and ensures the correct polarization and differentiation of the RPTE cells. In addition, ClC-5 facilitates acidification of the endolysosomal pathway, allowing degradation of bad-quality collagens. **(B)** In absence of ClC-5, cellular dedifferentiation is induced, possibly beginning by a decrease on MUC1/E-cadherin plasma membrane levels, which will release β-catenin that will be translocated to the nucleus and promote Col I/IV transcription. Consequently, collagen synthesis is increased and, together with impaired lysosomal degradation due to acidification defects, ends in massive intracellular accumulation of Col I /IV. Next, cells will steadily release these collagens to the external medium, also increasing their extracellular levels. Besides, due to the loss of cell polarity, Col I/IV are missecreted leading to abnormal accumulation that may lead to renal fibrosis over time. Abbreviations: e-Cadh.: e-Cadherin, ECM: extracellular matrix, ER: Endoplasmic Reticulum.

## MATERIALS AND METHODS

### Cell culture

RPTEC/TERT1 (#CRL-4031, ATCC®) cells were grown in Dulbecco’s Modified Eagle Medium: Nutrient Mixture F-12 (1:1, v/v) (#31331093, Thermo Fisher Scientific) supplemented with 20 mM HEPES (#15630–080, Gibco), 60 nM sodium selenite (#S9133, Sigma-Aldrich), 5 μg/ml transferring (#T1428, Sigma-Aldrich), 50 nM dexamethasone (#D8893, Sigma-Aldrich), 100 U/ml penicillin and 100 μg/ml streptomycin (#15240–062, Gibco), 2% foetal bovine serum (#10270, Gibco), 5 μg/ml insulin (#I9278, Sigma-Aldrich), 10 ng/ml epidermal growth factor (#E4127, Sigma-Aldrich) and 3 nM triiodothyronine (#T5516, Sigma-Aldrich). Culture conditions were maintained at 37°C in a humidified atmosphere containing 5% CO^2^. RPTEC/TERT1 control cells, *CLCN5* knockdown cells (*CLCN5* KD), and knockdown cells overexpressing ClC-5 WT form (ClC-5 rWT) were obtained by stable transfection as described in (18).

### Mycoplasma test

Mycoplasma contamination was analysed by PCR. The media of RPTEC/TERT1 cell line were collected and centrifuged at 300g for 3 min to remove cell debris. The supernatant was centrifuged at 15.000g for 10 min to sediment the mycoplasma. The supernatant was carefully discarded and re-suspended the pellet in milli-q water. Samples were heated at 95ºC for 5 minutes. Then, PCR amplification of Mycoplasma DNA was performed as described in Duran et al. Briefly, two μl of DNA was amplified in a 25 μl final volume of reaction mix containing 2.5 mM MgCl2, 0.2 mM dNTPs, 0.2 μM forward primer mix (5′-CGCCTGAGTAGTACGTWCGC -3′, 5′-TGCCTGRGTAGTACATTCGC-3’, 5’-CRCCTGAGTAGTATGCTCGC-3’, 5-’CGCCTGGGTAGTACATTCGC-3’) 0.2 μM reverse primer mix (5′-GCGGTGTGTACAARACCCGA-3′, 5’-GCGGTGTGTACAAACCCCGA -3’), and 1 unit of Taq polymerase (Bioline, Meridian Bioscience). The PCR conditions were an initial melting step of 94°C for 4 min; then 35 cycles of 94°C for 30 s, an annealing step of 55°C for 30 s, and 72°C for 45 s; followed by a final elongation step of 72°C for 6 min. PCR products were separated in 2% agarose gels.

### Clcn5^+/+^, Clcn5^+/-^ and Clcn5^-/-^ mice

Paraffined kidneys from *Clcn5*^*+/+*^, *Clcn5*^*+/-*^ and *Clcn5*^*-/-*^ mice (N=3 female mice for each condition, 3 month-old) were gently provided by Dr. Baisong Lu (Wake Forest Institute for Regenerative Medicine, USA) (20).

### H/E, Sirus Red/Fast Green and Col IV staining

Kidney sections were stained with H/E, Sirius Red/Fast Green Collagen or Col IV for histological analysis as described previously (21). Sirius Red was used to stain all types of collagens, whereas Fast Green was used to stain non-collagenous proteins. H/E-stained sections were evaluated by light microscopy using a semiquantitative analysis assessing the following parameters: epithelial hyperplasia (N, normal; 1, minimal; 2, mild; 3, moderate; 4, marked), inflammatory infiltrate in the lamina propria (N, normal; 1, minimal; 2, mild; 3, moderate; 4, marked), oedema of the lamina propria (0, normal; 1, minimal; 2, mild; 3, moderate; 4, marked), structure glomeruli (N, normal; A, altered), structure tubules (N, normal; A, altered). The histological study was performed in a blinded fashion.

### Measurement of basement membrane thickness

The thickness of the basement membrane was measured in kidney tissues sections using the ruler tool of the QuPath software (22). Ten different measures were performed per stain (Sirius Red/Fast Green and Col IV) of each mouse.

### Collagen secretion assay and western blot

The media of RPTEC/TERT1 cells was replaced with fresh medium for up 24 h to performed collagen secretion assay. The media were collected and centrifuged at 4000 rpm to removed cell debris. The supernatants (SN) were denatured at 95 °C for 5 min with Laemmli SDS sample buffer. For cell extracts, cells were washed with PBS 1x and lysed with SET 1x Buffer (10 mM Tris–HCl pH 7.4, 150 mM NaCl, 1 mM EDTA and 1% SDS). The cell extracts and SN were resolved in 15% SDS-PAGE gels and transferred to PVDF membranes (#ISEQ00010, Millipore) at 100V during 3 h. Membranes were blocked with non-fat dry milk diluted in PBS-T (1x PBS, 0.1% Tween-20) for 1 h and incubated overnight at 4°C with Collagen 1 (#ab34710, Abcam), Collagen IV (#ab6586, Abcam), MUC-1 (MA1-06503, Themofisher), E-Cadherin (610181, BD Transduction Labs), LAMP-1 (#53-1079-42), ClC-5 (#GTX53963, Gentex), and tubulin (T4026, Sigma) antibodies. Band intensities were visualized using Odyssey Fc Imaging System (Li-Cor).

### Collagen degradation assay

For collagen degradation assay, RPTEC/TERT1 cells were cultured into plates for 10 days to allow cell differentiation. Intracellular levels of collagen I and IV and SN after treatment with vehicle (DMSO), proteasome inhibitor (MG132; # 474790, Sigma)), bafilomycin A1(#19-148, Sigma) or lysosomal enzymatic activity inhibitor (Leupeptin; # L2884, Sigma) were analysed by WB as described previously.

### Cycloheximide chase assay

The turnover rates of Col I and IV was determined by cycloheximide (#C7698, Sigma)) chase assays. RPTEC/TERT1 cells were incubated with 50 μg/ml CHX or DMSO (vehicle) and collected at different times (0, 3, 6, 8, 24h).Then, cellular extracts were analysed by WB as describe previously.

### Microscopy and immunofluorescence

RPTEC/TERT1 cells were cultured on coverslips during 7 days. Cells were fixed with 4% paraformaldehyde for 20 min at room temperature. Aldehyde groups were quenched in 50 mM NH^4^Cl/PBS for 30 min and non-specific binding sites were blocked with 5% BSA in PBS for 60 min. Primary antibodies were diluted in blocking reagent and incubated overnight at 4 °C (Collagen I (# ab34710, Abcam, Collagen IV (#ab6586, Abcam), MUC-1 (MA1-06503, Themofisher), LAMP-1 (#53-1079-42), LAMP-1 (#53-1079-72, Invitrogen), KDEL(#Ab10C3, Abcam),, Beta-catenin (#ab32572, Abcam). Secondary antibodies conjugated with 488 (#A28175, Invitrogen) and 568 (#A28175, Invitrogen) were diluted in blocking solution and incubated for 1h at room temperature. Cells were incubated with Hoechst 33342 (1:2000 dilution, #H1399, Invitrogen) or Phalloidin (1:200 dilution, #R415, Invitrogen) for 1h at room temperature. Fluorescence was visualized in a confocal spectral Zeiss LSM 980 microscope. Images were processing using FIJI (23). Manders’ overlap coefficient was calculated using ImageJ plugin JACoP (24).

To determine the number, volume and area of LAMP1, Col I and Col IV positive elements, we used the 3D objects counter v2.0 tool from FIJI (24). All images for each experiment were taken on the same day under the same conditions and the same z-step (0.5 μm). The threshold was automatically set for each image. DAPI was used to count the number of nuclei per field.

### Real time quantitative RT-PCR

For gene expression analysis, total RNA was isolated from RPTEC/TERT1 cells using Trizol reagent following the manufacturer’s instructions. Equal amounts of RNA (1 μg) were retro transcribed to cDNA using the High Capacity RNA-to-cDNA Kit according to manufacturer’s instructions. Gene expression was evaluated by real-time quantitative PCR using SYBR green MasterMix (#A25742, Applied Biosystems) or TaqMan MasterMix (#4369016, Applied Biosystems), according to the manufacturer’s instructions. Total levels of *CLCN5*, Collagen I and IV, SOX 2 and 4, EFNB1 and TBP was evaluated with specific primers (*CLCN5* primers: 5′-GGGATAGGCACCGAGAGAT-3′ and 5′-GGTTAAACCAGAATCCCCCTGT-3′; Collagen I primers: 5’-GTGGTCAGGCTGGTGTGATG-3’ and 5’-CAGGGAGACCCTGGAATCCG-3’); Collagen IV primers: 5’-ATGGGGCCCCGGCTCAGC and 5’-ATCCTCTTTCACCTTTCAATAGC; SOX 2 primers: 5’-GAGCTTTGCAGGAAGTTTGC-3’ and 5’-GCAAGAAGCCTCTCCTTGAA-3’: SOX 9 primers: 5’-GTACCCGCACTTGCACAAC-3’ and 5’-CGCTCTCGTTCAGAAGTCTC-3’; EFNB1 primers: 5’-GTATCCTGGAGCTCCCTCAACC-3’ and 5’-GCTTGTAGTACTCATAGGGCCG-3’: TBP 5′-CGGCTGTTTAACTTCGCTTC-3′ and 5′-CAGACGCCAAGAAACAGTGA-3′). In contrast, totals levels of BMP4 were measured with a commercial Taqman probe (Hs00370078_m1). Data were normalized with TBP (Hs00427620_m1).Amplification protocol was performed using the LightCycler® 480 System (Roche). Relative expression fold change was determined by the comparative 2^(−ΔΔCT)^ method after normalizing to TBP.

### Endolysosomal pH determination

Endosomal pH was determined by using LysoSensor™ Yellow/blue DND-160 (Thermofisher, L7545). RPTEC/TERT1 cells were cultured in IBIDI® chambers for 10 days. Cells were washed 3 times with isotonic solution containing: 2.5 mM KCl, 140 mM NaCl, 1.2 mM CaCl_2_, 0.5 mM MgCl_2_, 5 mM Glucose and 10 mM HEPES (305 mosmol/l, pH 7.4 adjusted with Tris); and then treated with LysoSensor™ for 10 min at 37°C. After incubation period, cells were washed again with isotonic solution and imaged using a Leica Thunder Imager 3D Cell Culture Microscope for three minutes without stimulation. A continuation, cells were treated with vehicle (DMSO) or 100 nM Bafilomycin A1 to get a normalization point. Images were acquired with a 4X objective on Leica Thunder Imager 3D Cell Culture Microscope and processed using ImageJ software.

### Beta-catenin nuclear translocation

Nuclear translocation of beta-catenin was analysed by immunofluorescence. Briefly, cell lines were cultured on IBIDI chambers during 10 days and, after this period, starved overnight and exposed to 1.5 ng/ml of TFGβ (#100-B-001, R&D Systems) for 6 and 24 hours. At the end of treatment, cells were washed and lysed or fixed as describe previously. Fluorescence was visualized in a confocal spectral Zeiss LSM 980 microscope. Images were processing and analysed using FIJI (23).

### Statistical analysis

All data are means ± SEM. In all cases a D’Agostino– Pearson omnibus normality test was performed before any hypothesis contrast test. Statistical analysis and graphics were performed using GraphPad Prism 6 (RRID:SCR_002798) or SigmaPlot 10 (RRID:SCR_003210) software. For data that followed normal distributions, we applied either Student’s t test or one-way analysis of variance (ANOVA) followed by Tukey’s post hoc test. For data that did not fit a normal distribution, we used Mann–Whitney’s unpaired t test and nonparametric ANOVA (Kruskal–Wallis) followed by Dunn’s post hoc test. Binary and categorical variables were analysed by Fisher’s exact test. Criteria for a significant statistical difference were: *p<0.05; **p<0.01, ***p<0.001.

## ACKNOWLEDGMENTS

We thank the patient advocacy group ASDENT (Asociación de la Enfermedad de Dent, www.asdent.org) for its continuous support. We also thank all members of the Renal Physiopathology Group for valuable discussions, especially C. Castells for technical support. Specific primers to detect mice *Col4a1* and *Hprt1* were kindly provided by Dr. Jacobs. Fluorescence microscopy was performed at the High technology unit (UAT) at the Vall d’Hebron Research Institute (VHIR). This work reflects only the authors’ views, and the EU Community is not liable for any use that may be made of the information contained therein.

## AUTHOR CONTRIBUTIONS

MD: Conceptualization, Investigation, Methodology, Formal analysis, Validation, Writing – review and editing. GA: Investigation, Writing – review and editing. CCE: Investigation, Writing – review and editing. LB: Investigation, Writing – review and editing. MS: Formal analysis, Writing – review and editing. AM: Conceptualization, Formal analysis, Funding acquisition, Resources, Supervision, Writing – review and editing. GCR: Conceptualization, Formal analysis, Investigation, Methodology, Validation, Funding acquisition, Supervision, Writing – original draft, Writing – review and editing. All authors read and approved the final manuscript.

## DATA SHARING STATEMENT

The authors confirm that the data supporting the findings of this study are available within the article or its supplementary materials. Correspondence and material requests should be addressed to Dr. Cantero-Recasens or Dr. Meseguer.

