## Supplementary material for "Renal Cl-/H+ antiporter ClC-5 regulates collagen production and release in Dent Disease models"

[#] Both authors should be considered as joint last authors

[*] Correspondence to

Dr. Anna Meseguer:
Dr. Gerard Cantero-Recasens:

### INDEX

#### Supplementary figures

#### SUPPLEMENTARY FIGURE 1

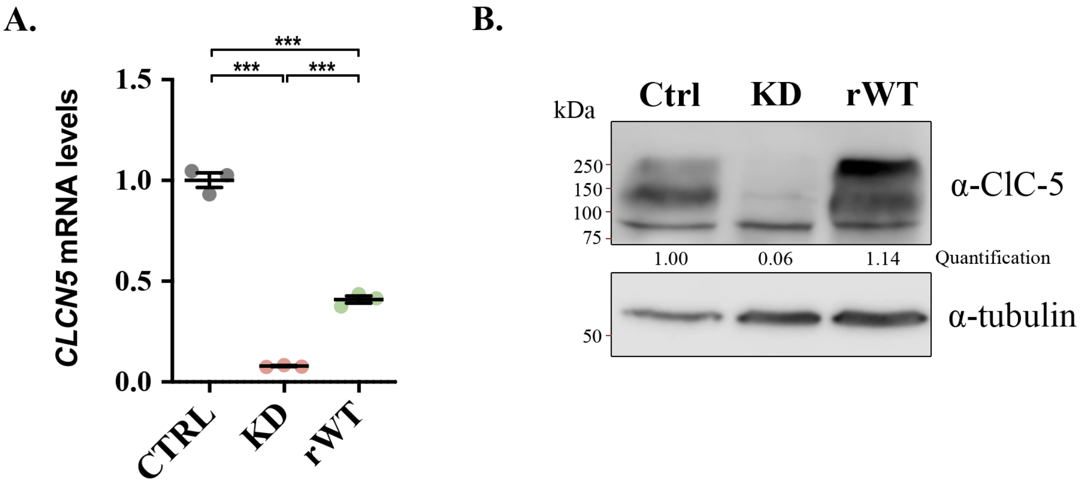

**Suppl. Figure 1.** Characterisation *CLCN5* KD and ClC-5 rWT cell lines. **(A)** *CLCN5* mRNA levels normalised to values of *TBP* from control (CTRL), *CLCN5* KD (KD) and ClC-5 rWT (rWT) cells. All values are represented as relative value compared to control cells. Average values ± SEM are plotted as scatter plot with bar graph. **(B)** Cell lysates from control (Ctrl), *CLCN5* KD (KD) and ClC-5 rWT (rWT) cells were analysed by western blot with an anti-ClC-5 to test expression levels. Tubulin was used as a loading control. Quantification of ClC-5 protein levels is provided. Abbreviations: Ctrl: Control, KD: *CLCN5* KD, rWT: ClC-5 rWT. Statistical significance was determined using one-way ANOVA followed by Tukey's post hoc test. *** $p<0.001$.

#### SUPPLEMENTARY FIGURE 2

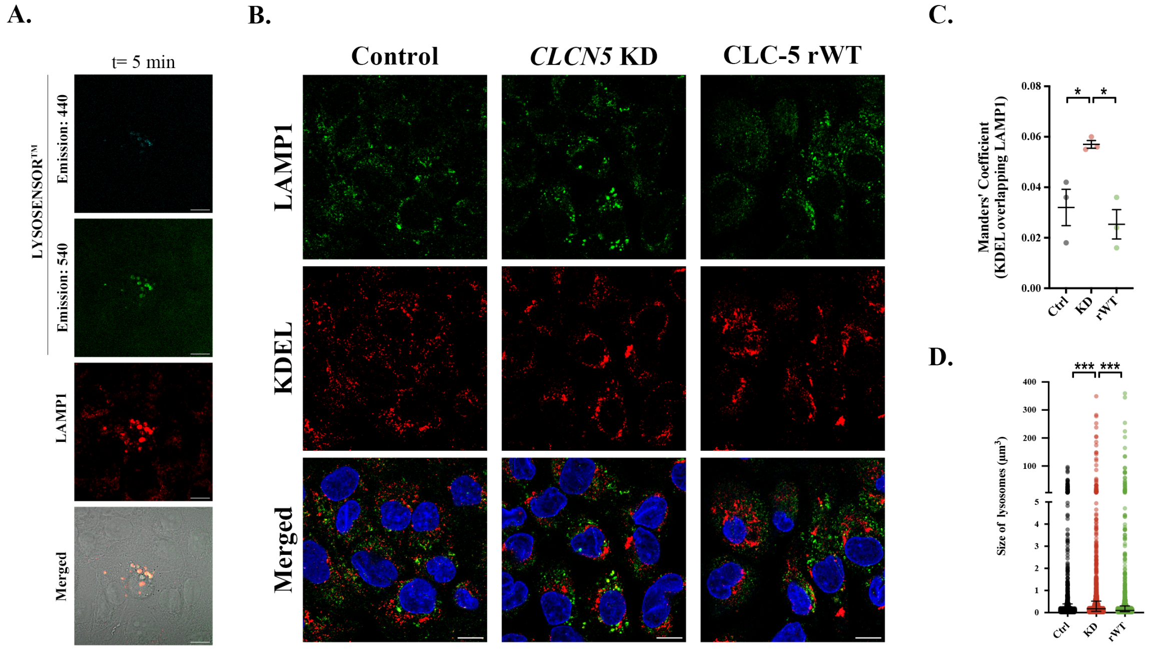

**Suppl. Figure 2.** Characterisation of the LysoSensor™ and lysosomes. **(A)** Control cells were incubated with the LysoSensor™ (Cyan, emission 440; Green, emission 540) for 5 min, processed for immunofluorescence and stained with an anti-LAMP1 (Red). Scale bars = 10 µm. Bright field is provided to confirm intracellular staining. **(B)** Representative immunofluorescence z-stack single-plane images of control, *CLCN5* KD, and ClC-5 rWT cells stained with anti-LAMP1 (Green), anti-KDEL (Red) and Hoechst 33342 (Blue). Scale bars = 10 µm. **(C)** Colocalization between LAMP1 and KDEL for different cell lines (Control, *CLCN5* KD and ClC-5 rWT) were calculated from immunofluorescence images by Manders' coefficient using FIJI. Average values ± SEM are plotted as scatter plot with bar graph. The y-axis represents Manders' coefficient of the fraction of KDEL overlapping with LAMP1. (D) Volume of lysosomes from control (black), *CLCN5* KD (KD, red) and ClC-5 rWT (rWT, green) cells was calculated from individual immunofluorescent images of LAMP1 using 3D analysis Fiji software. The y axis represents the volume of the lysosomes in cubic micrometres. Abbreviations: Ctrl: Control, KD: *CLCN5* KD, rWT: ClC-5 rWT. Statistical significance was determined using one-way ANOVA followed by Tukey's post hoc test. * $p<0.05$, ** $p<0.01$, *** $p<0.001$.

SUPPLEMENTARY FIGURE 3

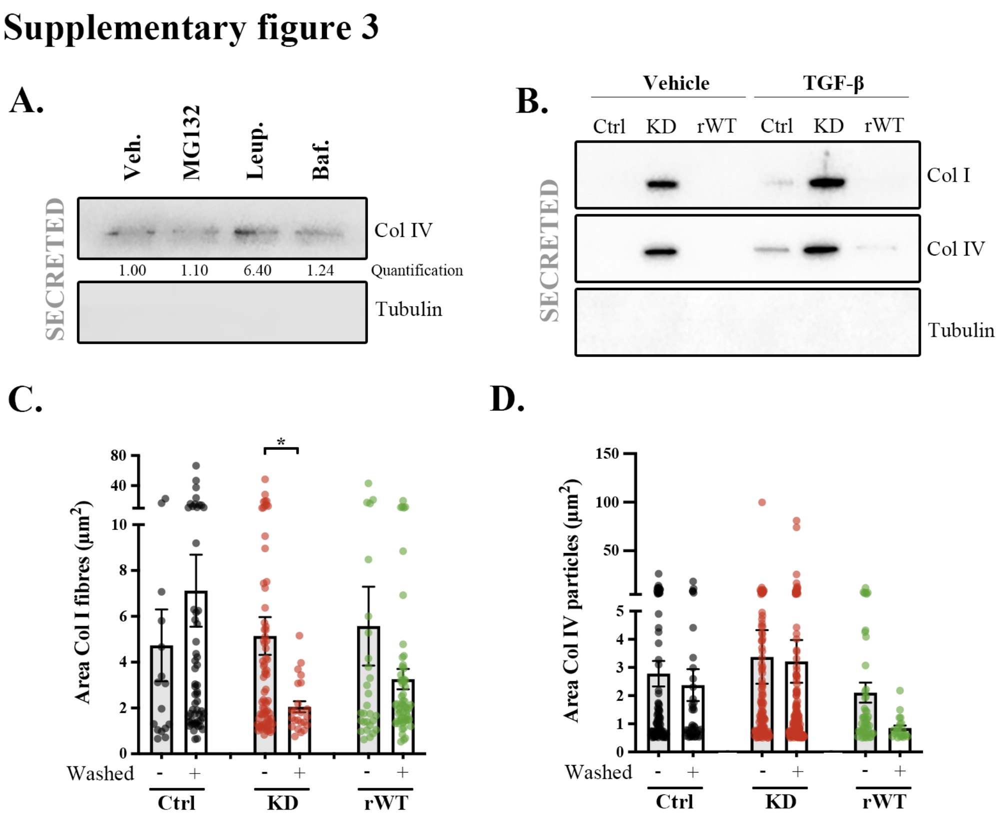

**Suppl. Figure 3.** Effect of *CLCN5* deletion on extracellular collagens. **(A)** Col IV extracellular (secreted) levels from control cells treated with vehicle (Veh.), MG132, Leupeptin (Leup.) or bafilomycin (Baf.). Tubulin was used as a loading control. Quantification of Col IV levels is included. **(B)** Col I and IV extracellular (secreted) levels from control (Ctrl), *CLCN5* KD (KD) and ClC-5 rWT (rWT) cells treated with vehicle or TGF-β. Tubulin was used as negative control of lysate contamination. **(C, D)** Area of Col I **(C)** or Col IV **(D)** fibres was calculated from immunofluorescence images of control, *CLCN5* KD and ClC-5 rWT cells before and after washing using FIJI software. The y axis represents the area of the fibres in square micrometres. Abbreviations: Ctrl: Control, KD: *CLCN5* KD, rWT: ClC-5 rWT. Statistical significance was determined using one-way ANOVA followed by Tukey's post hoc test. * p<0.05, ** p<0.01, *** p<0.001.

#### SUPPLEMENTARY TABLE 1

| Genotype | Haematoxylin & eosin (H/E) staining | | | | |
|---|---|---|---|---|---|
| | Epithelial hyperplasia | Inflammatory infiltrate | Oedema | Structure glomeruli | Structure tubules |
| **Clcn5$^{+/+}$** | N | N | 0 | N | N |
| **Clcn5$^{+/-}$** | N | N | 0 - 1 | N | N |
| **Clcn5$^{-/-}$** | N | N | 0 - 1 | N | N |

N: Normal
Score Oedema: 0: normal, 1: minimal
Evaluation of haematoxylin & eosin staining from 3 mice per genotype.

**Suppl. Table 1.** H/E staining's histological analysis of *Clcn5*$^{+/+}$, *Clcn5*$^{+/-}$ and *Clcn5*$^{-/-}$ mice. Evaluation of haematoxylin & eosin staining from 3 mice per genotype.

**SUPPLEMENTARY TABLE 2**

| Gene | Protein | Regulation | logFC | Adj P Val |
|---|---|---|---|---|
| **WNT7A** | Protein Wnt-7a | UP | 1.9 | 3.15E-12 |
| **SOX2** | Transcription factor SOX-2 | UP | 1.2 | 7.34E-08 |
| **EPHB2** | Ephrin type-B receptors | UP | 1.0 | 1.23E-10 |
| **WNT7B** | Protein Wnt-7b | UP | 1.0 | 2.77E-07 |
| **EDN1** | endothelin-1 | UP | 0.9 | 3.98E-05 |
| **SOX17** | Transcription factor SOX-17 | UP | 0.9 | 8.81E-07 |
| **WNT5A** | Protein Wnt-5a | UP | 0.8 | 8.92E-04 |
| **ENC1** | ectodermal-neural cortex 1 | UP | 0.8 | 6.92E-06 |
| **FGF18** | Fibroblast growth factor 18 | UP | 0.7 | 7.59E-04 |
| **TNFRSF11B** | Tumor necrosis factor receptor superfamily member 11B / Osteoprotegerin | UP | 0.7 | 9.57E-04 |
| **LBH** | Protein LBH | UP | 0.6 | 2.94E-03 |
| **ITF2 (TCF4)** | Immunoglobulin transcription factor 2 | UP | 0.6 | 3.67E-03 |
| **BMP4** | Bone morphogenetic protein 4 | UP | 0.6 | 1.05E-03 |
| **CCND1** | Cyclin-D1 | UP | 0.6 | 1.40E-03 |
| **PTTG** | Securin | UP | 0.5 | 4.53E-02 |
| **MYC** | Myc proto-oncogene protein | UP | 0.5 | 7.23E-03 |
| **CDKN2A** | p16INK4a | UP | 0.4 | 4.98E-03 |
| **FZD7** | Frizzled 7 | UP | 0.4 | 7.56E-03 |
| **FGF9** | Fibroblast growth factor 9 | UP | 0.3 | 3.18E-02 |
| **RUNX2** | Runt-related transcription factor 2 | UP | 0.3 | 3.46E-02 |
| **SOX9** | Transcription factor Sox-9 | DOWN | -0.5 | 1.75E-02 |
| **EFNB1** | Ephrin-B1 | DOWN | -1.4 | 1.26E-09 |

**Suppl. Table 2.** β-catenin target genes affected by *CLCN5* KD. Data extracted from Duran et al, 2021 (https://doi.org/10.1093/hmg/ddab131).
